# Effects of nitrous oxide and ketamine on the prefrontal cortex in mice: a comparative study

**DOI:** 10.1101/2022.09.19.508563

**Authors:** Stanislav Rozov, Roosa Saarreharju, Stanislav Khirug, Markus Storvik, Claudio Rivera, Tomi Rantamäki

## Abstract

Nitrous oxide (N_2_O; laughing gas) has recently been reported as a putative rapid-acting antidepressant, but little is known about the underlying mechanisms. We performed transcriptomics, *in situ* hybridization, and electrophysiological studies to examine the potential shared signatures induced by 1 h inhalation of 50% N_2_O and a single subanesthetic dose of ketamine in the medial prefrontal cortex (mPFC) in adult mice. Both treatments similarly affected the transcription of several negative regulators of mitogen-activated protein kinases (MAPKs), namely, dual specificity phosphatases. The effects were primarily located in the pyramidal cells. Notably, the overall effects of N_2_O on mRNA expression were much more prominent and widespread compared to ketamine. Ketamine caused an elevation of the spiking frequency of putative pyramidal neurons and increased gamma activity (30–100 Hz) of cortical local field potentials. However, N_2_O produced no such effects. Spiking amplitudes and spike-to-local field potential phase locking of putative pyramidal neurons and interneurons in this brain area showed no uniform changes across treatments. Thus, this study characterized the electrophysiological and transcriptome-wide changes in mPFC triggered by exposure to N_2_O and compared them with those caused by the rapid-acting antidepressant ketamine in terms of both the direction of their regulation and localization.

## Introduction

Nitrous oxide (N_2_O; laughing gas) has been used for several decades as an adjunct anesthetic, analgesic, and anxiolytic. Recently, N_2_O has shown promise as a rapid-acting antidepressant [1,2], likely owing to its antagonistic effects on *N*-methyl-D-aspartate receptors (NMDARs), similar to those of ketamine. The antidepressant properties of subanesthetic-dose ketamine are proposed to be associated with stimulatory effects on neural ensembles in cortical microcircuits that may occur through the blockade of NMDARs in inhibitory interneurons and subsequent disinhibition of glutamatergic synapses and/or inhibition of pre-synaptic NMDARs on excitatory synapses of pyramidal cortical neurons. Consequently, an increase in excitatory synapse drive is observed [3] following the modulation of molecular changes such as regulation of TrkB neurotrophin receptor and mitogen-activated protein kinase (MAPK) signaling [4]. Knowledge of the mechanisms underlying the antidepressant effects of N_2_O remains limited. However, our recent animal studies indicate that N_2_O and ketamine trigger similar time-dependent effects on TrkB and MAPK signaling in the cortex. Specifically, both treatments acutely increase MAPK phosphorylation, which is followed by a brain state characterized by slow-wave electroencephalographic (EEG) activity, during which MAPK phosphorylation decreases, whereas TrkB receptor signaling increases [5].

The objective of this study was to investigate electrophysiological and molecular responses of the prefrontal cortex of adult mice to clinically relevant doses of N_2_O and compare the results to those acquired after treatment with a subanesthetic dose of ketamine. Single unit activity (SUA) and phase coupling analyses showed non-uniform responses to treatment in putative pyramidal neurons and interneurons. Gene ontology and rank-rank hypergeometric overlap (RRHO) analyses of the whole-transcriptome sequencing data revealed that both treatments acutely affected the MAPK pathway, in particular the expression of dual-specificity phosphatases (DUSPs). Taken together, these findings demonstrate the presence of multiple molecular targets for N_2_O, some of which are also common to ketamine in terms of both direction of regulation and localization, which support our recent findings [4]. The heterogeneity of electrophysiological responses to both treatments suggests that the relationships within and between local circuits are more complex than proposed by the disinhibition hypothesis.

## Results

### Electrophysiological effects of treatments with ketamine and N_2_O

To characterize the acute electrophysiological responses to 50% N_2_O and subanesthetic-dose ketamine (10 mg/kg, i.p.), we first examined their effects on local field potentials (LFPs), firing rate, amplitude, and spike-to-LFP coupling of SUA in the anterior cingulate cortex (ACC; **Figure 1C**). Ketamine (**Figure 1A**) caused an elevation of gamma activity (30–100 Hz; perm. T-test; *p* < 0.05) in this cortical region, whereas N_2_O (**Figure 1B**) significantly increased theta rhythm (7–9 Hz) and suppressed beta activity (15–29 Hz; perm. T-test; *p* < 0.05). Spike sorting identified two major extracellular electrophysiological responses—which are characteristic of putative pyramidal neuron and interneuron cell types—on the basis of spike shapes and autocorrelograms (**Figure 1D, E**). Ketamine treatment increased the firing rate of approximately 70% of putative pyramidal SUA (**Figure 1F**; p = 0.002, paired T-test, **Supplementary table 2**), whereas N_2_O treatment increased the firing rate of only 38% of putative pyramidal SUA (**Figure 1G**; *p* = 0.76, paired T-test; **Supplementary table 2**). Moreover, approximately 60% of interneuron-like SUA in the ketamine-treated group (**Figure 1H**; *p* = 0.22, paired T-test) and 65% in the N_2_O-treated group (**Figure 1I**; *p* = 0.25, paired T-test) showed higher firing rates. No significant changes in spike amplitude were registered for both putative neuronal types (**Figure 1F-I**, lower panels).

**Figure 1.**
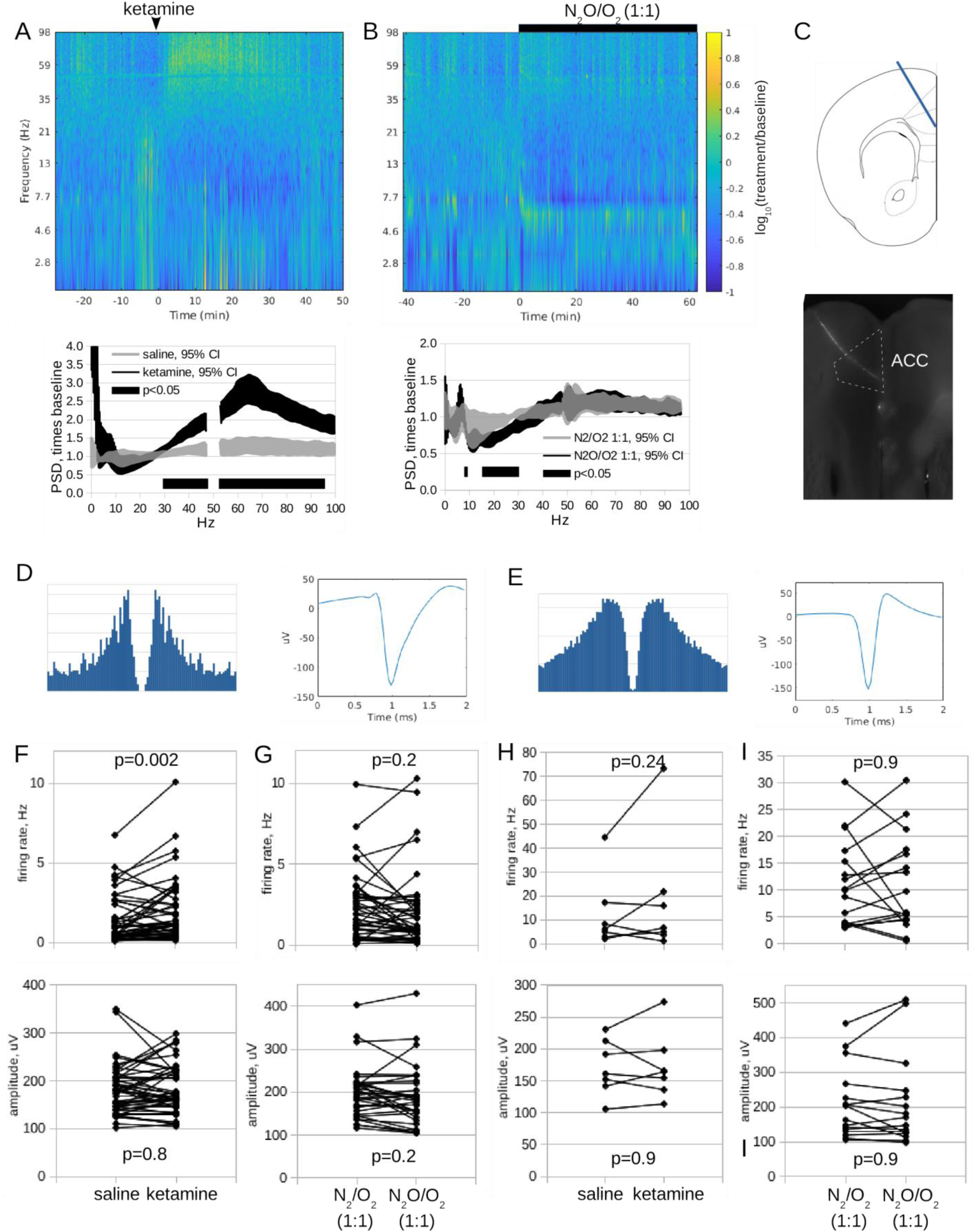
Effects of ketamine (10 mg/kg, i.p,) or 50% N_2_O/50% O_2_ on local field potentials and firing rates of putative pyramidal neurons and interneurons of the anterior cingulate cortex (ACC). **A**) Gamma activity (30– 100 Hz) considerably elevated in response to ketamine (spectrogram [upper panel]—ketamine administration is shown with an arrow head; normalized power spectrum [lower panel; black bars represent differences at *p* < 0.05; gap in the power spectrum at 50 Hz—mains interference removed by a notch filter]). **B**) N_2_O significantly increased theta rhythm (7–9 Hz) while suppressing beta activity (15–29 Hz) (spectrogram [upper panel]—black bar represents time of administration; normalized power spectrum [lower panel]). **C**) Topographical localization of probes in the ACC (upper panel) and its visualization with fluorescent tracer DiI (lower panel). Putative pyramidal neurons (**D**) and interneurons (**E**) were identified on the basis of their autocorrelograms (left panels) and spike shapes (right panels). Firing rates (upper panels) and amplitudes (lower panels) of putative pyramidal cells in response to saline and ketamine (**F**) or 50% N_2_/O_2_ and 50% N_2_O/O_2_ (**G**). Firing rates (upper panels) and amplitudes (lower panels) of putative interneurons in response to saline and ketamine (**H**) or 50% N_2_/O_2_ and 50% N_2_O/O_2_ (**I**). The number of subjects was 7 (N_2_O) and 6 (ketamine); total number of units was 44 + 18 (N_2_O) and 46 + 6 (ketamine).

Spectral analysis of the vicinity of spikes revealed dominant 3 Hz frequency and, in some cases, 7 Hz oscillations. Subsequent spike-to-LFP phase analysis followed by an omnibus test (histogram of spike-to-LFP phase distribution; **Supplementary figure 1E**) showed that approximately half of putative pyramidal SUA either coupled to the dominant 3 Hz rhythm irrespective of treatment with ketamine (**Supplementary figure 1A**; *p* = 0.83, Watson-Williams test) or N_2_O (**Supplementary figure 1B**; *p* = 0.76, Watson-Williams test) or developed coupling to that frequency in response to treatment. The other half of the SUA did not demonstrate coupling to any LFP frequencies (**Supplementary table 2**). In turn, virtually all interneuron-like SUA showed strong coupling to the dominant 3 Hz frequency, which was nevertheless not altered by either ketamine (**Supplementary figure 1C**; *p* = 0.56, Watson-Williams test) or N_2_O (**Supplementary figure 1D**; *p* = 0.56, Watson-Williams test); however, the strength of phase locking was somewhat reduced for both putative pyramidal neurons and interneurons in the case of N_2_O treatment (**Supplementary figure 1B,D**).

To examine changes in transcriptome induced by 50% N_2_O (60 min inhalation) or ketamine (10 mg/kg, i.p.), we conducted transcriptome-wide analysis of samples derived from mPFC 2 h after treatment onset (**Figure 2A**). Gene-wise comparison of groups treated with air or 50% N_2_O (15257 transcripts) revealed approximately 740 differentially expressed genes (Wald’s test, *p_adj_* < 0.05; **Figure 2A**). In contrast, out of 16968 transcripts analyzed in groups treated with saline or ketamine, only 15 were found to be differentially expressed: *Dusp5*, *Dusp6*, *Erf*, *Etv6*, *Fam84b*, *Ier5l*, *Junb*, *Kdm6b*, *Klf10*, *Lrtm2*, *Nr4a1*, *Sox8*, *Spry4*, *Tiparp*, and *Wisp1* (Wald’s test, *p_adj_* < 0.05; **Figure 2B)**.

**Figure 2.**
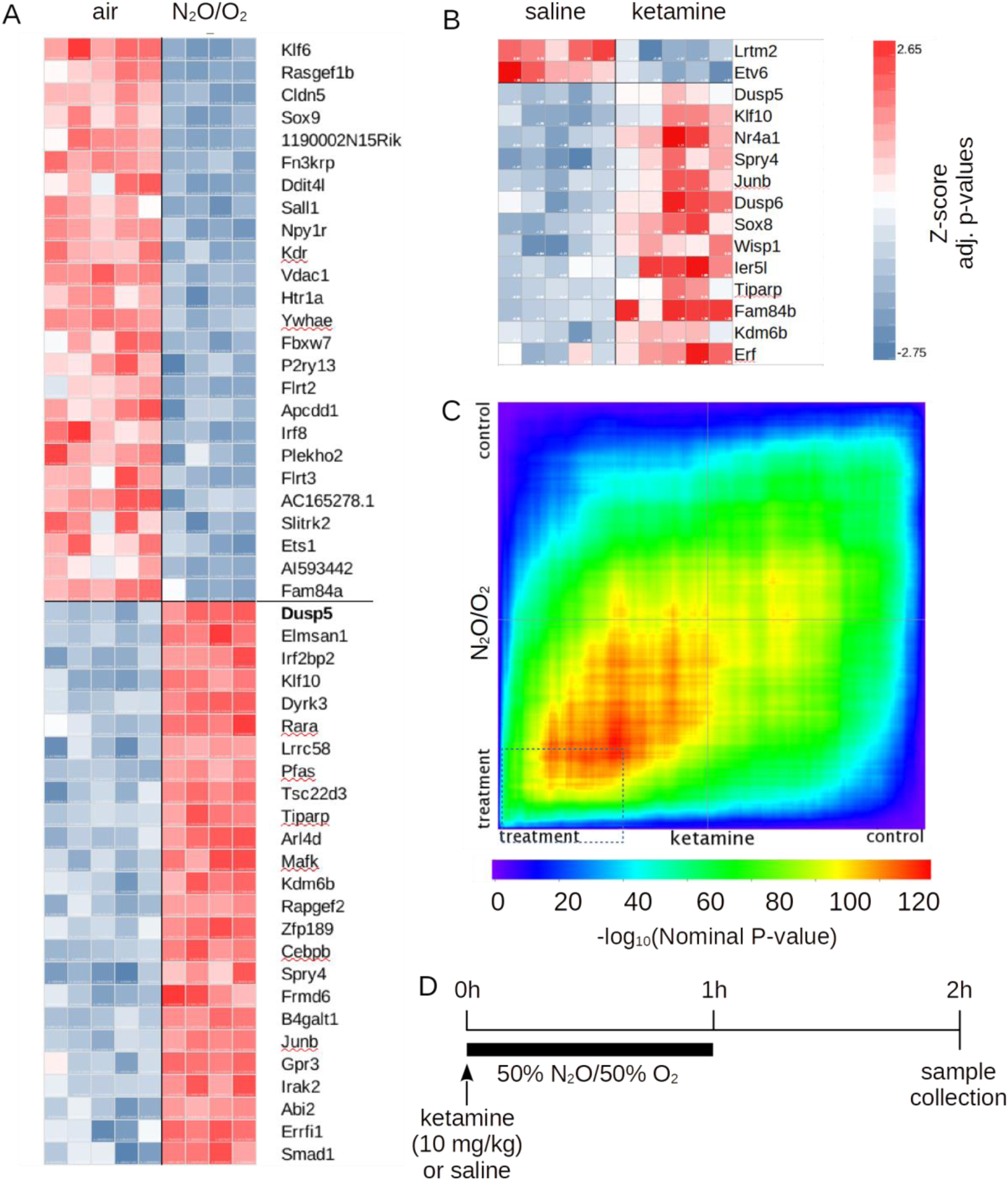
Results of RNAseq analysis of N_2_O- and ketamine-treated groups of animals in comparison with control treatment. Heatmap representation of expression levels of the 25 most significantly upregulated and downregulated genes between control and N_2_O (**A**) and control and ketamine (**B**) groups. Normalized count values are represented as Z scores; sorting based on adjusted p values. (**C**) Rank-rank hypergeometric overlap (RRHO) map representing shared mRNA changes in adult mouse medial prefrontal cortex (mPFC) in response to control vs. ketamine and control vs. N_2_O treatments. The thin dashed line represents the highest estimated probability of co-occurrence between these treatments (*p* = -log_10_128). (**D**) Schematic overview of the experiment: Male C57BL/6J mice were administered with saline (*n* = 5), ketamine (10 mg/kg, i.p., *n* = 5), or 50% N_2_O (60 min, *n* = 4). Two hours after the beginning of administration, samples were collected from the mPFC for further analysis.

A substantial portion of the pathways and gene transcripts affected by N_2_O are associated with synaptic function and neuronal activity (**Table 1**). These include synaptic vesicle trafficking factors (*Apba1*, *Cplx1*, *Stxbp2*, *Syt13*, *Snap25*, *Unc13a*, *Picalm*, and *Dnm3*), G-protein-coupled receptors (*Adgrl1*, *Chrm4*, *Gpr3*, *Htr1a*, *Htr2a*, *Htr6*, *Lpar1*, *Mc4r*, and *Ptger4*), and GABA_A_ receptor subunits (*Gabra2* and *Gabra3*). Furthermore, N_2_O downregulated the expression of postsynaptic scaffold factors such as *Nlgn2*, Nlgn3, and *Shank3* and affected the expression of several immediate early genes including *Arc*, *Egr1*, *Egr2*, *Egr4*, *Fos*, *Fosb*, *Junb*, *Klf4*, *Maff*, and *Nr4a1*. Notably, four of the 10 known dual-specificity serine-threonine phosphatases (DUSPs), which are key negative regulators of the three main pathways of the MAPK signaling cascade—JNK (*Dusp1*, *Dusp4*), p38-MAPK (*Dusp1*) and MAPK (*Dusp5*, *Dusp6*, *Dusp4*)—were regulated by N_2_O.

**Table 1.** Results of gene ontology analysis of genes, which expression level in response to treatment with N_2_O/O_2_ and/or ketamine significantly changed compared to control (Wald’s test, p<0.05).

| Biological Process (PANTHER GO-Slim) | over/under | Fold enrichment | raw P-value | FDR |
| --- | --- | --- | --- | --- |
| **N<sub>2</sub>O/O<sub>2</sub> (50%/50%, 60 min)** |  |  |  |  |
| epidermal cell differentiation (GO:0009913) | + | 16.22 | < 0.001 | 0.007 |
| circadian regulation of gene expression (GO:0032922) | + | 12.16 | < 0.001 | 0.001 |
| 'de novo' protein folding (GO:0006458) | + | 7.21 | 0.004 | 0.045 |
| transforming growth factor beta receptor signaling pathway (GO:0007179) | + | 6.28 | 0.001 | 0.014 |
| cellular response to peptide hormone stimulus (GO:0071375) | + | 5.16 | 0.001 | 0.014 |
| synaptic vesicle cycle (GO:0099504) | + | 4.42 | < 0.001 | 0.009 |
| autophagosome assembly (GO:0000045) | + | 4.32 | 0.004 | 0.046 |
| response to abiotic stimulus (GO:0009628) | + | 3.96 | < 0.001 | 0.005 |
| heart development (GO:0007507) | + | 3.72 | 0.004 | 0.047 |
| central nervous system development (GO:0007417) | + | 3.37 | 0.004 | 0.045 |
| MAPK cascade (GO:0000165) | + | 3.09 | < 0.001 | 0.008 |
| transmembrane receptor protein serine/threonine kinase signaling pathway (GO:0007178) | + | 3.07 | 0.004 | 0.046 |
| neuron projection morphogenesis (GO:0048812) | + | 2.57 | 0.002 | 0.027 |
| G protein-coupled receptor signaling pathway, coupled to cyclic nucleotide second messenger (GO:0007187) | + | 2.48 | 0.004 | 0.042 |
| chemical synaptic transmission (GO:0007268) | + | 2.36 | 0.001 | 0.017 |
| positive regulation of cellular protein metabolic process (GO:0032270) | + | 2.01 | 0.004 | 0.047 |
| regulation of transcription, DNA-templated (GO:0006355) | + | 1.72 | < 0.001 | < 0.001 |
| immune response (GO:0006955) | - | 0.44 | 0.003 | 0.039 |
| **Ketamine (10 mg/kg, i.p.)** |  |  |  |  |
| tissue morphogenesis (GO:0048729) | + | 84,9 | < 0.001 | 0.0193 |
| MAPK cascade (GO:0000165) | + | 32,05 | < 0.001 | 0.0123 |
| **N<sub>2</sub>O/O<sub>2</sub> and ketamine (RRHO analysis)** |  |  |  |  |
| MAPK cascade (GO:0000165) | + | 10.5 | < 0.001 | < 0.001 |
| regulation of transcription, DNA-templated (GO:0006355) | + | 2.48 | < 0.001 | < 0.001 |
| <u>apoptotic process (GO: 0006915)</u> | + | 6,72 | < 0.001 | 0.018 |
| <u>regulation of cell cycle (GO: 0051726)</u> | + | 5.54 | < 0.001 | 0.0027 |

Next, we used RRHO analysis to further analyze the concordance/discordance in mRNA changes in the mPFC induced by N_2_O and ketamine. Non-corrected results of Wald’s tests from the previous step were subjected to RRHO analysis, which revealed ∼1600 genes with the highest estimated probability of co-occurrence between these treatments (**Figure 2C**, *p* = -log_10_128). A subset of these genes, chosen based on *a posteriori* power analysis (alpha = 0.1, beta = 0.25, 125 genes), were further processed for GO search. This analysis showed 18 biological processes affected by N_2_O and only two processes regulated by ketamine (**Table 1**). Moreover, several genes that were significantly influenced by both treatments are known members of the MAPK family as well as a subset of immediate early genes (**Table 1**).

To validate the RNAseq findings, we performed qPCR using selected transcripts. The expression levels of *Dusp1*, *Dusp5*, *Dusp6*, and *Nr4a1* were significantly increased by both ketamine and N_2_O treatment, whereas only N_2_O treatment increased the expression of *Fos* and *Arc* (**Supplementary figure 2A**). Notably, in these experiments, we compared N_2_O-treated group with a non-treated (air) control group, which raises questions about the potential effects of 50% O_2_ on target genes. To address this concern, we compared the expression of *Dusp1*, *Dusp5*, and *Dusp*6 in the 50% N2O/ 50% O_2_ group and 50% N_2_/50% O_2_ treatment and found no differences between the treatments (**Supplementary figure 2A-B**).

The results of transcriptome analysis prompted us to examine the spatial distribution of *Fos*, *Arc*, *Nr4a1*, *Dusp1,* and *Dusp6* transcripts and study their co-localization with markers of glutamatergic (*Slc17a7*) and GABAergic (*Gad2*) neurons. Triple fluorescent *in situ* hybridization chain reaction (FISHCR) revealed cortical layers 2-3 of mPFC as a primary place of expression of transcripts of interest (**Figure 3 A-E, upper panels**). In addition, *Arc* was expressed in the inner cortical layers (presumably 5-6), and expression of *Nr4a1* was relatively evenly spread across the cortex in the case of saline and ketamine administration. Notably, *Fos*, *Arc*, *Nr4a1*, *Dusp1,* and *Dusp6* transcripts were primarily (M = 0.65–0.85) co-localized with *Slc17a7* (marker of excitatory synapses; **Figure 3F** and **Supplementary figure 3**) and only marginally (*M* = 0.05–0.20) with *Gad2* (marker of GABAergic neurons). This distribution and co-localization was observed regardless of treatment (*p* > 0.05; non-paired T-test).

**Figure 3.**
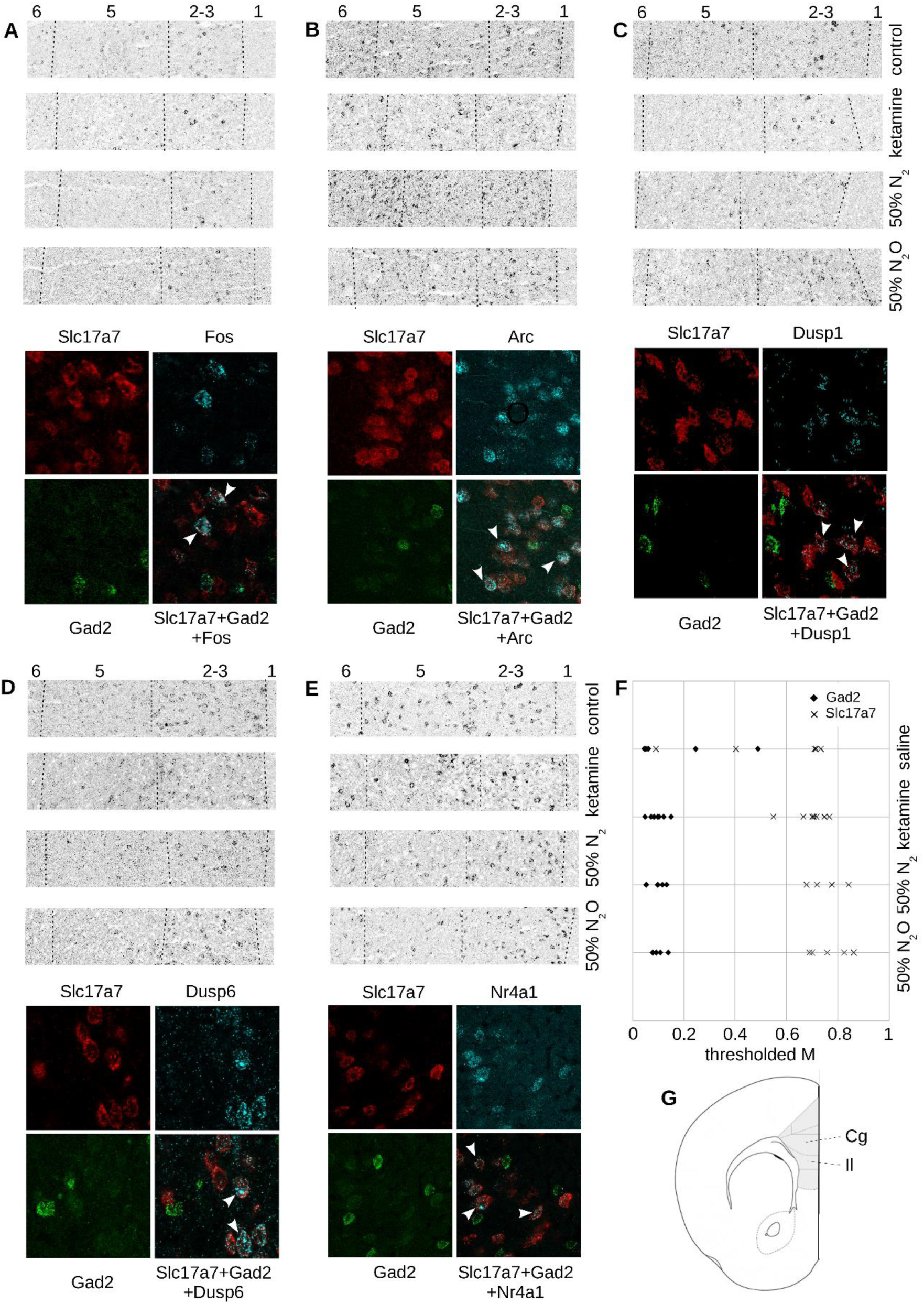
Co-localization of *Fos*, *Arc*, *Dusp1*, *Dusp6*, and *Nr4a1* transcripts with markers of glutamatergic (*Slc17a7*) and GABAergic (*Gad2*) neurons was assessed using fluorescent *in situ* hybridization chain reaction. Fragments of the medial prefrontal cortex (mPFC) labeled with oligonucleotide probe sets against *Fos* (A), *Arc* (B), *Dusp1* (C), *Dusp6* (D), and *Nr4a1* (E) (upper panels; fluorescence is pseudo-colored in grayscale) and their colocalization (cyan) with *Slc17a7* (red) and *Gad2* (green) markers (lower panels; fluorescence is pseudo-colored in red, green, and cyan). Dashed lines represent approximate borders between cortical layers; arrowheads point to cells with overlapped signals. F) Fraction of Fos-positive cells co-localized with *Slc17a7*-or *Gad2*-positive cells in the mPFC, as measured by Mander’s coefficient. G) Region of interest used for co-localization analysis (shadowed area) and a part of the mPFC used in the upper panels of A–E (rectangular outline).

## Discussion

N_2_O has shown promise as a putative rapid-acting antidepressant [1] and has several favorable translational properties compared to ketamine. N_2_O is administered via inhalation, does not undergo biotransformation [25], and shows relatively similar (rapid) pharmacokinetics across mammalian species. Similar to ketamine, N_2_O acts as an NMDAR antagonist. However, the precise mechanisms underlying the antidepressant effects of N_2_O remain poorly studied. Our comparative RNA screening analysis of the mouse mPFC, a brain area recognized for its role in the development of depressive states in humans and animal models of depression [26], suggests that the MAPK signaling pathway is a common target of both N_2_O and subanesthetic-dose ketamine. In particular, both treatments upregulated the expression of *Dusp*s, negative regulators of MAPK signaling. This cascade has been consistently associated with the antidepressant effects of ketamine [27], and both MAPK and DUSPs are also activated by non-pharmacological treatments of depression such as electroconvulsive shock (a model of electroconvulsive therapy) [28]. Previous transcriptomic studies addressing the effects of ketamine have reported varying results [29–31]. However, the MAPK pathway has been highlighted in one of these studies [31].

Recently, we demonstrated N_2_O-mediated elevation of *Dusp1* (*MKP1;* [5]), however, no consensus has been reached concerning the effects of ketamine on this target, as it has been shown to be either upregulated [29] or downregulated [31] in the prefrontal cortex of rodents. In contrast, MAPK pathway (ERK1/2) inhibitors have been shown to cause depressive-like behavior [32] and the phosphorylation of MAPKs has been demonstrated to be altered in depressed patients and in animal models of depression [33]. Moreover, inhibition of ERK1/2 activity blocks antidepressant-like behavioral effects of antidepressant drugs, including ketamine [34].

The ability of ketamine to regulate MAPKs is likely associated with its stimulatory effects on neural ensembles in cortical microcircuits [35]. These effects are suggested to arise through the blockade of NMDARs in either inhibitory interneurons and subsequent disinhibition of glutamatergic synapses or direct inhibition of pre-synaptic NMDARs on excitatory synapses of pyramidal cortical neurons, which leads to an increase in excitatory synapse drive [3]. Our *in situ* data show that the expression of *Dusp1* and *Dusp6* was primarily co-localized with markers of glutamatergic cells, which may either support the latter hypothesis or indicate that the activation of these genes was not directly linked to the inhibition of NMDARs in interneurons. N_2_O-induced activation of *Dusps* may be related to either its ability to block NMDARs [36] or its interaction with serotonergic [37] and noradrenergic [38] receptors and cation channels [39].

The diversity of targets of N_2_O suggests complex pharmacological effects of this drug. Indeed, treatment with N2O affected the transcription of not only several genes associated with the MAPK pathway but also hundreds of other genes, including synaptic vesicle trafficking factors (*Apba1*, *Cplx1*, *Stxbp2*, *Syt13*, *Snap25*, *Unc13a*, *Picalm*, *Dnm3*), G-protein-coupled receptors (*Adgrl1*, *Chrm4*, *Gpr3*, *Htr1a*, *Htr2a*, *Htr6*, *Lpar1*, *Mc4r* and *Ptger4*), ion channels (*Gabra2* and *Gabra3*), postsynaptic scaffold factors (*Nlgn2*, *Nlgn3*, *Shank3),* and immediate early genes (*Arc*, *Egr1*, *Egr2*, *Fos Egr4*, *Fosb*, *Junb*, *Klf4*, *Maff* and *Nr4a1),* which may be regulated by neuronal activity and subsequent activation of MAPK (35). Notably, effects of ketamine were much less prominent. However, one of the hits, *Nr4a1* (also known as *Nurr77*), has also previously been shown to be upregulated by ketamine in primary neuronal cultures [40].

Previous studies have shown that *Dusps* and *Nr4a1* were upregulated in animal models of depression; however, the results vary depending on the brain region and method used to generate the model [32,41,42]. Expression of *Arc* was downregulated in the frontal cortex and CA1 area of the hippocampus both after chronic mild stress and in a genetic animal model of depression [43]. Effects of N_2_O have been rarely studied in animals [44]; therefore, assessment of the expression of these targets in a model subjected to N_2_O treatment would provide a better understanding of its putative antidepressant properties.

Our study demonstrated an increase in high-frequency (>30 Hz) cortical LFPs in response to ketamine treatment, which is consistent with several reports in humans and rodents [5,45]. Strong enhancement of the 7 Hz band and suppression of beta activity (15–30 Hz) were induced by N_2_O. To bridge these macroscopic EEG observations to SUA recorded, we assessed SUA in relation to LFPs registered at the same location and time. Spike-to-LFP coupling analysis showed that although a 3 Hz oscillation was dominant in the vicinity of spikes in all analyzed SUA, units that demonstrated persistent coupling to this oscillation were neither synchronous nor changed in response to treatment with either compound. Additionally, a small fraction of SUA had a second dominant frequency at 7 Hz, which is likely related to the increase in its power observed in spectrograms. Another noteworthy observation that is yet to be explained is the stronger coupling of putative interneurons to the 3 Hz activity than that of pyramidal cells. Slow oscillations during wakefulness in frontal cortical areas have been attributed to the activity of several circuits. Recent reports [46] show strong phase-locked spiking activity of orbitofrontal neurons in response to the 4 Hz rhythm of respiration. However, similar to our finding, Kőszeghy et al. [47] demonstrated modest phase locking of mPFC neurons to this rhythm with no phase preference throughout their population. Another explanation may be synchronization of the cortico-amygdala circuit, which is related to fear response [48], or the cortico-ventral tegmental area-hypothalamic circuit, which may be related to working memory [49].

Furthermore, we observed a significant increase in firing rate in the majority of recorded putative pyramidal cells in response to ketamine, which was consistent with the observed elevation of gamma activity and is in line with recent findings [50]; moreover, in combination with our *in situ* data showing primary colocalization of immediate early genes *Fos* and *Arc* with vesicular glutamate transporter 1, the finding aligns with the inhibition-disinhibition hypothesis [3]. However, in addition to these cells, a fraction of interneuron-like SUA also showed increased spiking frequency, whereas the firing rate of the rest of them decreased moderately. In contrast, no increase in the firing rate of putative pyramidal cells was observed in the N_2_O-treated group. Because of the small number of recorded responses from putative interneurons, which indicates the overall low relative numbers of these cells in the cerebral cortex, it is not possible to generalize observed effects. Nevertheless, our data show heterogeneity in the response of both cell types to ketamine and marked difference in the firing rate response between both treatment groups, suggesting more complex relationships within and between local circuits than a simple on-off response. Recording with higher electrode density and quantity accompanied by optogenetic stimulation/inhibition of selected cell types would not only provide certainty regarding cell types under investigation, including types of interneurons, but also prove essential to revealing changes in connectivity of local cortical circuits in response to treatments.

Collectively, our study characterized transcriptome-wide changes in mPFC triggered by exposure to N_2_O and compared them with those caused by the rapid-acting antidepressant ketamine in terms of both the direction of their regulation and localization, thus further supporting our recent observations [4]. In addition, the heterogeneity of electrophysiological responses to both treatments suggests more complex relationships within and between local circuits than proposed by the disinhibition hypothesis.

## Materials and Methods

### Animals

Approximately 8–10-week-old, male C57BL/6JOlaHsd mice (Envigo, Netherlands) were used. Animals were housed singly in sound-proof cages (Scantainer, Scanbur, Sweden) under controlled conditions (22 ±1°C, 12 h light-dark cycle, 6 AM lights on) in the animal facility of the University of Helsinki with free access to food and water, unless specified otherwise. Treatments and behavioral tests, excluding the saccharin preference test, were performed during the light phase (*Zeitgeber Time*, ZT 3-7). The experiments were performed according to the guidelines of the Society for Neuroscience and were approved by the County Administrative Board of Southern Finland (License: ESAVI/5844/2019).

### Drug treatments

Ketamine-HCl (Ketaminol®, Intervet International B.V., 511485) was diluted in saline and injected intraperitoneally (i.p.) at a dose of 10 mg/kg [6] (injection volume ∼10 ml/kg). Controls received saline. A 1:1 mixture of N_2_O and O_2_ (Woikoski, Finland) was administered for 40–60 min, as described in previous clinical [1] and animal [5] studies. Because the RNAseq experiment included one control group for both ketamine and N_2_O treatments, saline (i.p.) was administered to both the control and N_2_O groups to control for the effects of injection. To control for the potential effects of hyperoxygenation in the 50% N_2_O treatment group, an additional experiment was performed using a 50% N_2_/50% O_2_ mixture.

### Collection of brain samples

For the biochemical analyses, the animals were euthanized by cervical dislocation 2 h after ketamine/saline injections or 2 h after the onset of 1 h N_2_O treatment. The whole brain or bilateral mPFC (including prelimbic and infralimbic cortices) tissue was rapidly dissected on a cooled dish [7] and stored at -80 °C until further processing.

### RNA sequencing

mRNA extraction was performed with the NucleoSpin RNA Plus (Macherey-Nagel, Germany) kit in accordance with the manufacturer’s instructions. Before sequencing, RNA integrity was tested with a Bioanalyzer (Agilent Technologies, CA, USA), and samples with RIN >9.0 were used in analysis. Samples were sequenced with the Illumina NextSeq sequencer (Illumina, San Diego, CA, USA) at the Biomedicum Functional Genomics Unit (FuGU) in High output runs using the NEBNext® Ultra™ II Directional RNA Library Prep Kit for Illumina 3. The sequencing was performed as single-end 75 sequencing (SE75 or 1×75bp; read length 75 bp).

Data processing steps included FastQC quality analysis, analysis summarization with MultiQC [8], light quality trimming with Trimmomatic [9], alignment of the sample reads against the mouse GRCm38.p6 (GCA_000001635.8) reference with spliced transcript alignment to a reference genome (STAR) [10], mapping quality assessment with Qualimap [11], read quantification (featureCounts, [12]), differential expression statistics (negative binomial linear model followed by Wald’s test, followed by Benjamini–Hochberg correction for multiple comparisons implemented in DESeq2 R library [13]. To identify the molecular processes primarily affected by ketamine or N_2_O alone, gene expression data (Wald’s test, *p_adj_* < 0.05) were subjected to gene ontology (GO, [14]) analysis, and pathways that exceeded the significance threshold (Fisher’s exact test, *p_adj_* < 0.05) were considered as significantly enriched. To identify the common molecular signatures of ketamine and N_2_O effects, signed log10(P) values from the previous step were used in rank-rank hypergeometric overlap (RRHO) analysis as described earlier (step size = 100) [15]. *A posteriori* power analysis with alpha = 0.1 and beta = 0.25 [16] was used as a selection criterion for a commonly regulated subset of genes identified by RRHO (signed log10(P) values higher than maximum estimated co-occurrence rate), to be included in the GO search as described above. Genes that were associated with several pathways and had a >1.5 times difference vs. control (Wald’s test, *p_adj_* < 0.05) were selected for the qPCR validation analysis.

### Quantitative RT-PCR

mRNA extraction was done with the NucleoSpin RNA Plus (Macherey-Nagel, Germany) kit according to the manufacturer’s instructions. RNA concentration and purity were assessed with the Nanodrop 2000 Spectrophotometer (Millipore, USA). Samples with an RNA concentration >100 ng/µl, 260/280 nm ratio >1.9, and 260/230 nm ratio > 1.9 were processed for cDNA synthesis (one sample excluded) using the Maxima First Strand cDNA Synthesis Kit and dsDNase mix (K1672, Thermo Scientific, USA). The primers used to amplify specific cDNA regions of the transcripts are listed in **Supplementary table 1**. Primers for *Dusp1*, *Dusp5*, *Dusp6*, and *Nr4a1* were designed using the PrimerBLAST service (NCBI, USA) and tested for specificity and selectivity. Amplification of cDNA and corresponding control samples was done with Maxima SYBR Green/ROX qPCR Master Mix (2X) (K0221, Thermo Scientific, USA) on the Lightcycler® 480 System (Roche, Switzerland). Relative quantification of templates was performed as described in [17], with cDNA data being normalized to the joint *Gapdh* and *beta-actin* transcript levels. Amplification efficiency of PCR reactions was estimated using a dilution series: *Gapdh*: 1.93; *Actb*:1.88, *Fos*: 1.84, *Nr4a1*: 1.91, *Arc*: 1.98, *Dusp1*:1.99, *Dusp5*: 2.04, *Dusp6*: 1.97.

### Fluorescent in situ hybridization chain reaction (FISHCR)

Sets of probes for *Fos* (accession # NM_010234.3), *Arc* (NM_018790.3), *Dusp1* (NM_013642.3), *Dusp6* (NM_026268.3), *Nr4a1* (NM_010444.3), *Slc17a7* (NM_182993), and *Gad2* (NM_008078.2) as well as buffers for hybridization, washing, and amplification steps were produced by and purchased from Molecular Instruments (USA). Twenty micrometer-thick coronal sections of mPFC were prepared at -15°C with the Leica CM1860 UV (Leica, Germany) and stored at -80°C until further processing. Prior to FISHCR, sections were fixed in cold 4% paraformaldehyde for 15 min and processed according to the supplier’s instructions [18] with the following modifications: concentrations of probes in hybridization solutions were 1 nM (*Slc17a7* and *Gad2*) and 16 nM (*Fos*, *Arc*, *Dusp1*, *Dusp6I,* and *Nr4a1*); *Fos*, *Arc*, *Dusp1*, *Dusp6*, and *Nr4a1* probes were designed to link to the B4 amplifier and were conjugated with Alexa488 dye, and *Slc17a7* and *Gad2* probes were conjugated with B2-Alexa647 and B3-Alexa546, respectively, to perform the triple hybridization reaction; and the incubation times for hybridization and amplification steps were 20 h. Sections were mounted in Fluoromount-G media (Thermo Fisher Scientific, USA). Data acquisition was performed with a Leica DMi8 system equipped with a HC PL APO 40x/1.25 motCORR objective, White Light Laser Stellaris 8, HyD S/HyD X detectors, and LASX software. Image background correction (BaSiC; [19]), stitching, thresholding, and co-localization analysis (BIOP JaCoP; [20]) were performed using Fiji software v. 1.53 [21].

### Electrophysiological recording and data analysis

Mice were anesthetized using isoflurane (5% induction, 2% maintenance), accompanied by the administration of analgesic (carprofen, s.c. 5 mg/kg) and local analgesic (lidocaine), and craniotomy for electrophysiological recording and montage of a metal frame for head fixation was performed. To facilitate the administration of injectable compounds during recording sessions, the subjects were additionally implanted with a biocompatible catheter (s.c.) connected to the injection port. Animals were allowed to recover for 7 days, followed by 4 days of habituation to Mobile HomeCage® (Neurotar, Helsinki, Finland). Recordings were performed in the anterior cingulate cortex (AP: - 2.0, ML: 0.75, angle 45°) on head-restrained conscious subjects with an A4×8-5mm-100-200-177-A32 probe (Neuronexus, USA) connected to an A32 preamplifier and SmartBox Pro amplifier (Neuronexus, USA). A face mask was used for the delivery of gaseous substances. Data acquisition was performed at a sampling rate of 20–30 kHz. At 40 min after control treatment, subjects received either ketamine or N_2_O, and recording continued for another 40 min. After the experiment, probe locations were verified on brain slices under a fluorescent microscope.

Power spectra were normalized using the frequency-wise division of power spectral density values of treatment phases per power spectral density values of baseline recordings. We used non-paired t-tests to compare normalized values between the control and treatment groups. To correct for multiple comparisons, we set a threshold for the vectors of t-values to retain only those values that exceeded *p* = 0.01, excluding the remainder from further analysis. Integrated t-values were computed for any contiguous array (cluster) of suprathreshold values. We tested the significance of these values against the distribution of integrated t-values of clusters acquired through the permutation-based generation of sets of matrices of t-values, followed by setting a threshold as described above. We considered clusters with *p* < 0.05 as significant.

Spike sorting was performed on band-pass filtered (300-5000 Hz, Butterworth filter, order 3), whitened, median-subtracted data using SpykingCircus software [22]. Results of spike sorting were manually verified using Phy2 software to identify putative pyramidal neuron- and interneuron-like spikes on the basis of the duration of the after-hyperpolarization phase and autocorrelogram analysis of inter-spike intervals. Spike-to-LFP coupling, firing rate assessment, and statistical analyses (omnibus test to assess the spike-to-LFP phase lock of single neurons, Watson-Williams test to assess the significance of treatment-induced spike-to-LFP phase lock change of multiple SUA, paired T-test to assess the significance of treatment-induced firing rate change) were performed in Matlab (Natick, USA) with the CircStat toolbox [23].

### Statistics

Statistical analysis of qPCR data was performed using R software (version 3.5.2, [24]). Differences among experimental groups in qPCR analysis were determined using either a two-tailed, non-paired T-test or one-way analysis of variance (ANOVA), followed by Tukey *post-hoc* test with Bonferroni correction.

## Acknowledgments

We are grateful to the personnel at the animal facility of the University of Helsinki for helping to conduct animal experiments and take care of the animals. We would like to thank the Mouse Behavioral Phenotyping Facility (MBPF) at the University of Helsinki for providing equipment and professional assistance for behavioral testing and mouse model implementation and the Functional Genomics Unit (FUGU), University of Helsinki, for assistance with RNAseq data analysis. We express gratitude to Woikoski OY for the provision of oxygen and nitrous oxide. Additionally, we thank Okko Alitalo and Dr. Samuel Kohtala for helping to conduct the experiments and improve the manuscript.

## Author Contributions

S.R., R.S., and T.R. planned the experiments; S.R. and R.S. performed the experiments; S.R. and R.S. prepared the figures; S.R. and R.S. performed statistical analysis; S.K. and C.R. assisted with electrophysiological experiments and corresponding data analysis; M.S. assisted with the pathway analyses and interpretation of the RNA sequencing data; S.R. and T.R. wrote the manuscript; and S.R., C.R., and T.R. acquired funding. All authors reviewed the manuscript and accepted the final submitted version.

---

**Supplementary figure 1.**
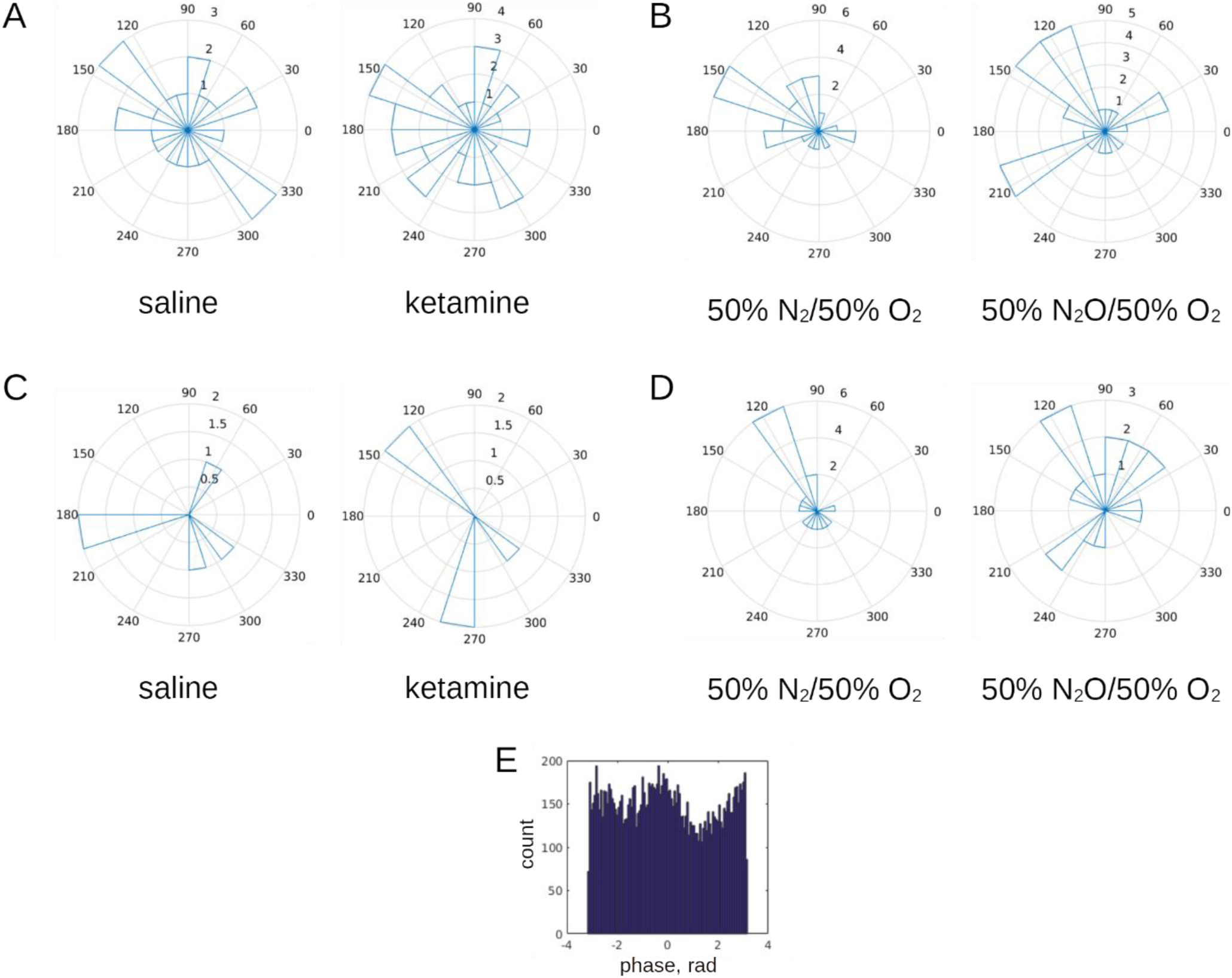
Effects of saline, ketamine (10 mg/kg, i.p,), 50% N_2_ / 50% O_2_ or 50% N_2_O / 50% O_2_ on spike-to-LFP phase coupling assessed at 3 Hz which was revealed as a dominant frequency in all examined SUA. A half of observed SUA of putative pyramidal cells had significant coupling to 3 Hz oscillation irrespectively of treatment (putative pyramidal cells: ketamine (**A**) and N_2_O (**B**); all SUA of putative interneurons were coupled to 3 Hz: ketamine (**C**) and N_2_O (**D**)). Example of spike-to-LFP phase coupling of putative pyramidal cell (**E**). The number of subjects was n=7 (N_2_O) and 6 (ketamine); total number of units was 44+18 (N_2_O) and 46+6 (ketamine).

**Supplementary figure 2.**
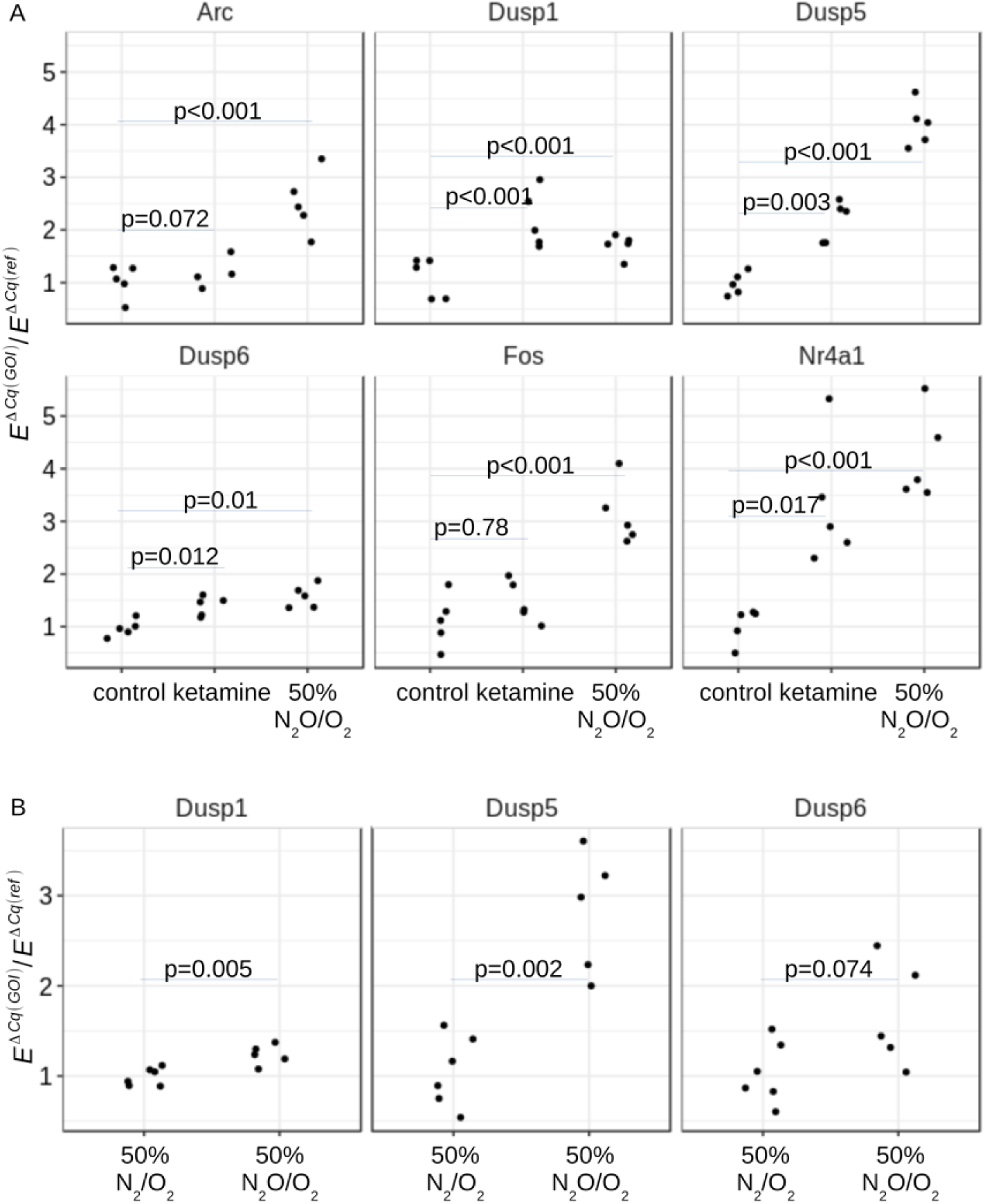
(**A**) Results of qPCR validation of expression of genes, selected based on RRHO analysis and fulfilled criteria of *a-posteriori* power test (alpha=0.1, beta=0.25). Data were analyzed gene-wise with two-tailed non-paired Student’s T-test (n=5). (**B**) Exposure to 50% O_2_ *per se* does not evoke any changes in *Dusp1,5* and *6*. Subjects were treated with either 50% N_2_O / 50% O_2_ or 50% N_2_ / 50% O_2_. Only group, treated with N_2_O showed significant alterations in *Dusp1* and *Dusp5* gene expression levels.

**Supplementary figure 3.**
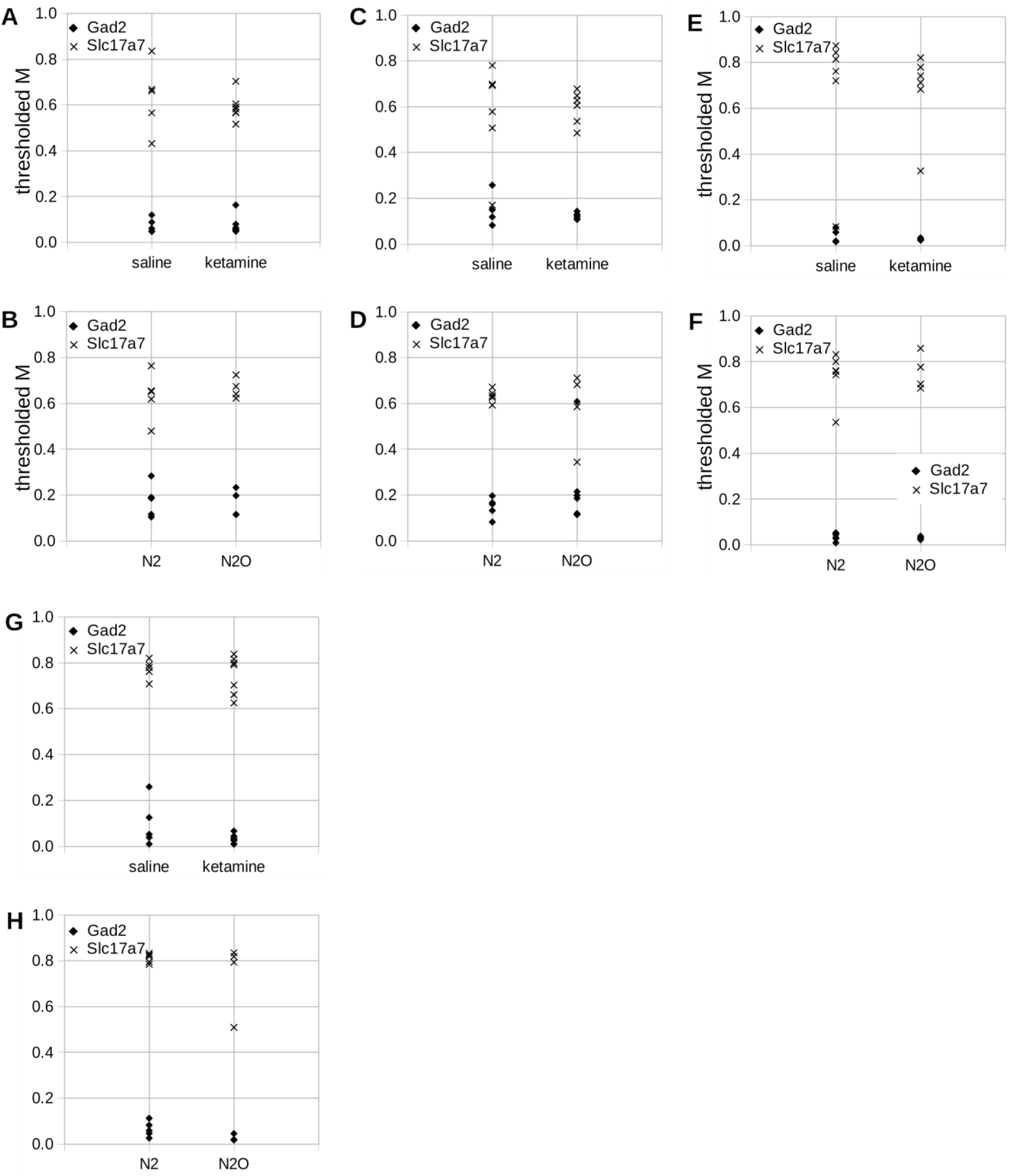
Fractions of *Arc*- (A,B), *Dusp1*- (C,D), *Dusp6*- (E,F), and *Nr4a1*- (G,H) positive cells co- localized with *Slc17a7*- or *Gad2*-positive cells in the mPFC as measured by Mander’s coefficient in saline and ketamine (A,C,E,G) or 50% 50% N_2_ / 50% O_2_ andN_2_O / 50% O_2_ (B,D,F,H).

**Supplementary table 1.**
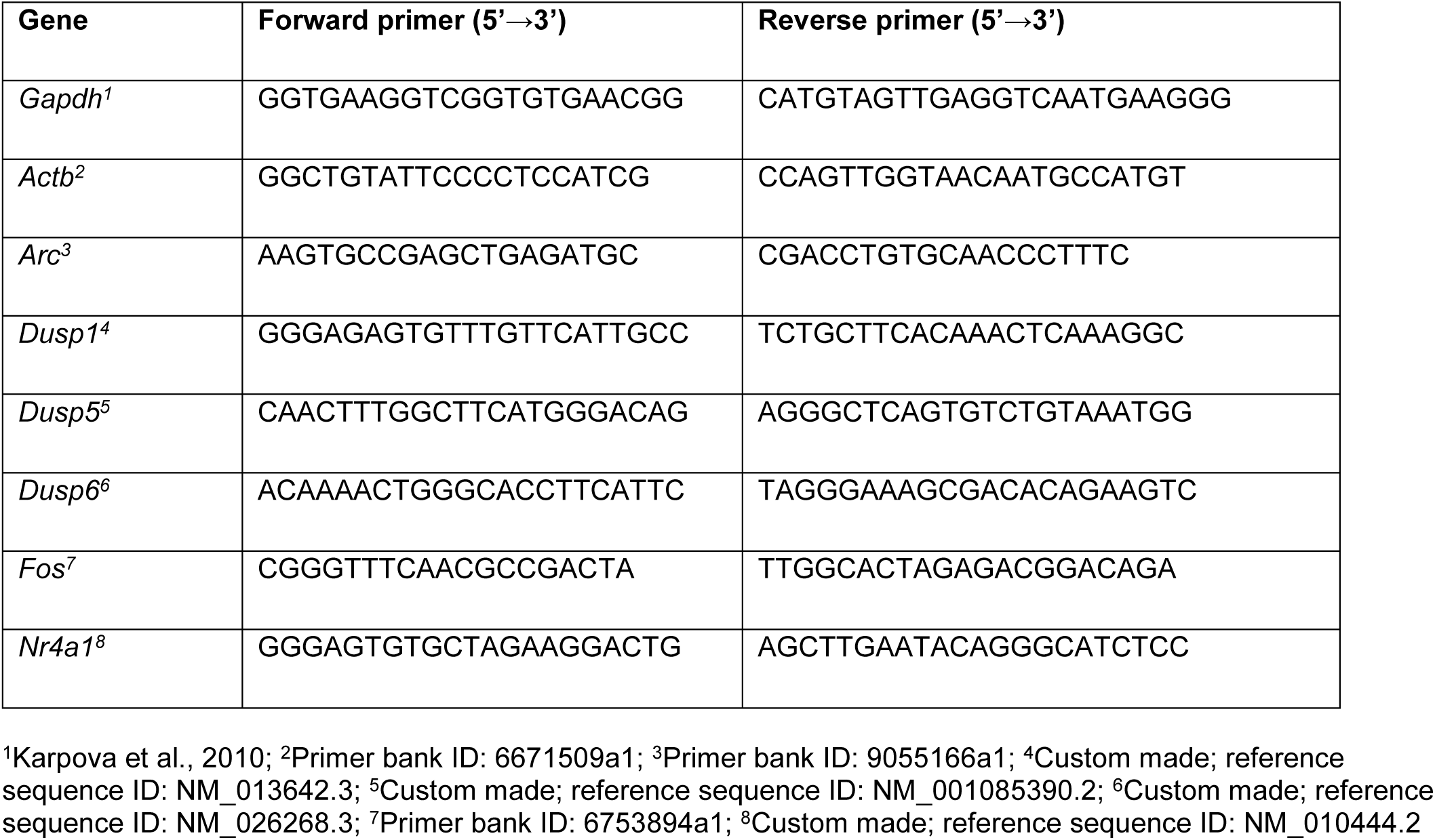
The primers used for quantitative polymerase chain reaction.

**Supplementary table 2.** Distribution of spike-to-LFP coupling status in response to treatment.

|  | ketamine | N<sub>2</sub>O |
| --- | --- | --- |
| SUA, putative pyramidal neurons, total | 46 | 44 |
| of those, with coupling lost after treatment | 1 | 7 |
| of those, with coupling appeared after treatment | 13 | 9 |
| of those, with no coupling | 9 | 8 |
| of those, with coupling persisted regardless of treatment | 23 | 20 |
| SUA, putative interneurons, total | 6 | 18 |
| of those, with coupling lost after treatment | 0 | 1 |
| of those, with coupling appeared after treatment | 0 | 0 |
| of those, with no coupling | 1 | 0 |
| of those, with coupling persisted regardless of treatment | 5 | 17 |

## References

1. Nagele P, Duma A, Kopec M, Gebara MA, Parsoei A, Walker M, et al. Nitrous Oxide for Treatment-Resistant Major Depression: A Proof-of-Concept Trial. Biol Psychiatry. 2015;78:10–18.

2. Nagele P, Palanca BJ, Gott B, Brown F, Barnes L, Nguyen T, et al. A phase 2 trial of inhaled nitrous oxide for treatment-resistant major depression. Science Translational Medicine. 2021;13:eabe1376.

3. Miller OH, Moran JT, Hall BJ. Two cellular hypotheses explaining the initiation of ketamine’s antidepressant actions: Direct inhibition and disinhibition. Neuropharmacology. 2016;100:17–26.

4. Kohtala S, Alitalo O, Rosenholm M, Rozov S, Rantamäki T. Time is of the essence: Coupling sleep-wake and circadian neurobiology to the antidepressant effects of ketamine. Pharmacol Ther. 2021;221:107741.

5. Kohtala S, Theilmann W, Rosenholm M, Penna L, Karabulut G, Uusitalo S, et al. Cortical Excitability and Activation of TrkB Signaling During Rebound Slow Oscillations Are Critical for Rapid Antidepressant Responses. Mol Neurobiol. 2019;56:4163–4174.

6. Moda-Sava RN, Murdock MH, Parekh PK, Fetcho RN, Huang BS, Huynh TN, et al. Sustained rescue of prefrontal circuit dysfunction by antidepressant-induced spine formation. Science. 2019;364:eaat8078.

7. Kohtala S, Theilmann W, Suomi T, Wigren H-K, Porkka-Heiskanen T, Elo LL, et al. Brief Isoflurane Anesthesia Produces Prominent Phosphoproteomic Changes in the Adult Mouse Hippocampus. ACS Chem Neurosci. 2016;7:749–756.

8. Ewels P, Magnusson M, Lundin S, Käller M. MultiQC: summarize analysis results for multiple tools and samples in a single report. Bioinformatics. 2016;32:3047–3048.

9. Bolger AM, Lohse M, Usadel B. Trimmomatic: a flexible trimmer for Illumina sequence data. Bioinformatics. 2014;30:2114–2120.

10. Dobin A, Davis CA, Schlesinger F, Drenkow J, Zaleski C, Jha S, et al. STAR: ultrafast universal RNA-seq aligner. Bioinformatics. 2013;29:15–21.

11. García-Alcalde F, Okonechnikov K, Carbonell J, Cruz LM, Götz S, Tarazona S, et al. Qualimap: evaluating next-generation sequencing alignment data. Bioinformatics. 2012;28:2678–2679.

12. Liao Y, Smyth GK, Shi W. featureCounts: an efficient general purpose program for assigning sequence reads to genomic features. Bioinformatics. 2014;30:923–930.

13. Love MI, Huber W, Anders S. Moderated estimation of fold change and dispersion for RNA-seq data with DESeq2. Genome Biol. 2014;15:550.

14. Mi H, Muruganujan A, Ebert D, Huang X, Thomas PD. PANTHER version 14: more genomes, a new PANTHER GO-slim and improvements in enrichment analysis tools. Nucleic Acids Res. 2019;47:D419–D426.

15. Plaisier SB, Taschereau R, Wong JA, Graeber TG. Rank-rank hypergeometric overlap: identification of statistically significant overlap between gene-expression signatures. Nucleic Acids Res. 2010;38:e169.

16. Hart SN, Therneau TM, Zhang Y, Poland GA, Kocher J-P. Calculating sample size estimates for RNA sequencing data. J Comput Biol. 2013;20:970–978.

17. Pfaffl MW. A new mathematical model for relative quantification in real-time RT–PCR. Nucleic Acids Res. 2001;29:e45.

18. Choi HMT, Schwarzkopf M, Fornace ME, Acharya A, Artavanis G, Stegmaier J, et al. Third-generation in situ hybridization chain reaction: multiplexed, quantitative, sensitive, versatile, robust. Development. 2018;145:dev165753.

19. Peng T, Thorn K, Schroeder T, Wang L, Theis FJ, Marr C, et al. A BaSiC tool for background and shading correction of optical microscopy images. Nat Commun. 2017;8:14836.

20. Bolte S, Cordelières FP. A guided tour into subcellular colocalization analysis in light microscopy. J Microsc. 2006;224:213–232.

21. Schneider CA, Rasband WS, Eliceiri KW. NIH Image to ImageJ: 25 years of image analysis. Nat Methods. 2012;9:671–675.

22. Yger P, Spampinato GL, Esposito E, Lefebvre B, Deny S, Gardella C, et al. A spike sorting toolbox for up to thousands of electrodes validated with ground truth recordings in vitro and in vivo. Elife. 2018;7:e34518.

23. Berens P. CircStat: A MATLAB Toolbox for Circular Statistics. Journal of Statistical Software. 2009;31:1–21.

24. R core team. R: A language and environment for statistical computing. R Foundation for Statistical Computing. 2018. 2018.

25. Banks A, Hardman JG. Nitrous oxide. Continuing Education in Anaesthesia Critical Care & Pain. 2005;5:145–148.

26. Wellman CL, Bollinger JL, Moench KM. Effects of stress on the structure and function of the medial prefrontal cortex: Insights from animal models. Int Rev Neurobiol. 2020;150:129–153.

27. Humo M, Ayazgök B, Becker LJ, Waltisperger E, Rantamäki T, Yalcin I. Ketamine induces rapid and sustained antidepressant-like effects in chronic pain induced depression: Role of MAPK signaling pathway. Prog Neuropsychopharmacol Biol Psychiatry. 2020;100:109898.

28. Kodama M, Russell DS, Duman RS. Electroconvulsive seizures increase the expression of MAP kinase phosphatases in limbic regions of rat brain. Neuropsychopharmacology. 2005;30:360–371.

29. Bagot RC, Cates HM, Purushothaman I, Vialou V, Heller EA, Yieh L, et al. Ketamine and Imipramine Reverse Transcriptional Signatures of Susceptibility and Induce Resilience-Specific Gene Expression Profiles. Biol Psychiatry. 2017;81:285–295.

30. Orozco-Solis R, Montellier E, Aguilar-Arnal L, Sato S, Vawter MP, Bunney BG, et al. A Circadian Genomic Signature Common to Ketamine and Sleep Deprivation in the Anterior Cingulate Cortex. Biol Psychiatry. 2017;82:351–360.

31. Ficek J, Zygmunt M, Piechota M, Hoinkis D, Rodriguez Parkitna J, Przewlocki R, et al. Molecular profile of dissociative drug ketamine in relation to its rapid antidepressant action. BMC Genomics. 2016;17:362.

32. Duric V, Banasr M, Licznerski P, Schmidt HD, Stockmeier CA, Simen AA, et al. A negative regulator of MAP kinase causes depressive behavior. Nat Med. 2010;16:1328–1332.

33. Meller E, Shen C, Nikolao TA, Jensen C, Tsimberg Y, Chen J, et al. Region-specific effects of acute and repeated restraint stress on the phosphorylation of mitogen-activated protein kinases. Brain Res. 2003;979:57– 64.

34. Duman CH, Schlesinger L, Kodama M, Russell DS, Duman RS. A role for MAP kinase signaling in behavioral models of depression and antidepressant treatment. Biol Psychiatry. 2007;61:661–670.

35. Rantamäki T, Kohtala S. Encoding, Consolidation, and Renormalization in Depression: Synaptic Homeostasis, Plasticity, and Sleep Integrate Rapid Antidepressant Effects. Pharmacol Rev. 2020;72:439–465.

36. Jevtović-Todorović V, Todorović SM, Mennerick S, Powell S, Dikranian K, Benshoff N, et al. Nitrous oxide (laughing gas) is an NMDA antagonist, neuroprotectant and neurotoxin. Nat Med. 1998;4:460–463.

37. Mukaida K, Shichino T, Fukuda K. Nitrous oxide increases serotonin release in the rat spinal cord. J Anesth. 2007;21:433–435.

38. Tyssowski KM, DeStefino NR, Cho J-H, Dunn CJ, Poston RG, Carty CE, et al. Different Neuronal Activity Patterns Induce Different Gene Expression Programs. Neuron. 2018;98:530–546.e11.

39. Gruss M, Bushell TJ, Bright DP, Lieb WR, Mathie A, Franks NP. Two-pore-domain K+ channels are a novel target for the anesthetic gases xenon, nitrous oxide, and cyclopropane. Mol Pharmacol. 2004;65:443–452.

40. Choi M, Lee SH, Wang SE, Ko SY, Song M, Choi J-S, et al. Ketamine produces antidepressant-like effects through phosphorylation-dependent nuclear export of histone deacetylase 5 (HDAC5) in rats. Proc Natl Acad Sci U S A. 2015;112:15755–15760.

41. Gourley SL, Wu FJ, Taylor JR. Corticosterone regulates pERK1/2 map kinase in a chronic depression model. Ann N Y Acad Sci. 2008;1148:509–514.

42. Jeanneteau F, Barrère C, Vos M, De Vries CJM, Rouillard C, Levesque D, et al. The Stress-Induced Transcription Factor NR4A1 Adjusts Mitochondrial Function and Synapse Number in Prefrontal Cortex. J Neurosci. 2018;38:1335–1350.

43. Elizalde N, Pastor PM, Garcia-García AL, Serres F, Venzala E, Huarte J, et al. Regulation of markers of synaptic function in mouse models of depression: chronic mild stress and decreased expression of VGLUT1. J Neurochem. 2010;114:1302–1314.

44. Liu W, Li Q, Ye B, Cao H, Shen F, Xu Z, et al. Repeated Nitrous Oxide Exposure Exerts Antidepressant-Like Effects Through Neuronal Nitric Oxide Synthase Activation in the Medial Prefrontal Cortex. Front Psychiatry. 2020;11:837.

45. Foster BL, Liley DTJ. Nitrous Oxide Paradoxically Modulates Slow Electroencephalogram Oscillations: Implications for Anesthesia Monitoring. Anesthesia & Analgesia. 2011;113:758–765.

46. Biskamp J, Bartos M, Sauer J-F. Organization of prefrontal network activity by respiration-related oscillations. Sci Rep. 2017;7:45508.

47. Kőszeghy Á, Lasztóczi B, Forro T, Klausberger T. Spike-Timing of Orbitofrontal Neurons Is Synchronized With Breathing. Front Cell Neurosci. 2018;12:105.

48. Karalis N, Dejean C, Chaudun F, Khoder S, Rozeske RR, Wurtz H, et al. 4 Hz oscillations synchronize prefrontal-amygdala circuits during fear behaviour. Nat Neurosci. 2016;19:605–612.

49. Fujisawa S, Buzsáki G. A 4 Hz oscillation adaptively synchronizes prefrontal, VTA, and hippocampal activities. Neuron. 2011;72:153–165.

50. Ali F, Gerhard DM, Sweasy K, Pothula S, Pittenger C, Duman RS, et al. Ketamine disinhibits dendrites and enhances calcium signals in prefrontal dendritic spines. Nat Commun. 2020;11:72.

